# A lethal human H5N5 influenza virus isolate exhibits low pandemic risk traits

**DOI:** 10.64898/2026.07.20.739507

**Authors:** Michelle N. Vu, Grace E. Quirk, Alexis E. Smathers, Kaitlyn Busfield-Thomason, Alaina L. Dorazio, Katelyn R. Domke, G. M. Humber, John Sembrat, Anita K. McElroy, Valerie Le Sage, Seema S. Lakdawala

## Abstract

In fall of 2025, a fatal infection of highly pathogenic avian influenza (HPAI) virus H5N5 occurred. To define the risk of this emerging virus to humans, we performed a comprehensive analysis based on our established triage. Serological analysis revealed that humans across all birth years had no detectable neutralizing antibodies to this H5N5 isolate. Further characterization revealed a lack of phenotypic signatures associated with epidemiologically successful influenza viruses in humans, including reduced replication in human airway cells and an avian-like pH of inactivation. Additionally, assessment of H5N5 in ferrets revealed a lack of direct contact transmission and moderate disease severity. H5N5 infection in ferrets with prior immunity against the 2009 H1N1 pandemic strain resulted in fewer clinical signs and reduced viral shedding. Together our data suggest that the current H5N5 HPAI lineage poses a low pandemic risk.

**Importance:** HPAI H5N5 viruses have caused widespread infection and death in avian species, and characterizing their pandemic risk traits is critical to understanding the threat posed to humans. In this work we analyzed an isolate that resulted in a human fatality in 2025. We found that this strain lacks many key features of influenza viruses with epidemiological success in humans including reduced growth in human lung cultures, a pH of inactivation less than 5.0, and lack of transmission to cohoused recipient ferrets. Prior immunity with seasonal H1N1 strain also reduced the viral load and disease burden of the virus. Taken together, these data suggest that currently circulating H5N5 poses a low risk to humans but highlights the importance of phenotypic characterizations for future risk assessments as the virus evolves in wild birds.

## Introduction

Influenza A viruses (IAV) have an incredibly broad host range, with wild birds serving as the natural reservoir for highly pathogenic avian influenza (HPAI) virus subtypes. HPAI clade 2.3.4.4b H5Nx viruses have caused widespread infection and death in avian species (1). Clade 2.3.4.4b is predominantly composed of H5N1 subtype viruses but has shown the propensity to reassort, to generate novel subtypes, including H5N5. HPAI subtype H5N5 virus was first detected in China around 2008 on poultry farms and live bird markets (2–4) and later in southwest Europe during the 2020-2021 season (5). Incursion into North America occurred via seabird migration with fatal detections in avian and mammalian wildlife in 2023 (6). Given its high prevalence in nature and the novel neuraminidase, H5N5 poses a risk to humans.

H5N5 caused its first human infection, which was fatal, in November 2025 in the US in Washington (7). The individual had underlying health conditions, and after contact with backyard ducks, developed severe symptoms including fever and respiratory distress. This H5N5 isolate contains a clade 2.3.4.4b hemagglutinin and internal gene segments cluster with currently circulating Eurasian lineage clade 6 (EA6) strains (7). Given the severity of H5N5 infections in poultry and the recent human spillover, we assessed the pandemic potential of this lethal human H5N5 isolate strain (A/Washington/2148/2025) *in vitro* and *in vivo* using our previously described pandemic risk triage pipeline (8).

## Results

### Absence of neutralizing antibodies against H5N5 in a representative human population

The human population has pre-existing IAV immunity, which may provide protection against emerging IAV strains. To assess antibody-mediated cross-protection against H5N5, serum samples obtained from healthy adults across a range of ages were tested for neutralizing antibodies. None of the individuals tested exhibited neutralizing antibodies against H5N5 above the limit of detection, whereas most individuals had neutralizing antibodies against the 2009 H1N1 pandemic strain (A/California/07/2009; herein referred to as H1N1pdm09) (Figure 1A), indicating that most adults lack neutralizing antibodies against H5N5 and are likely susceptible to H5N5 infection.

**Figure 1.**
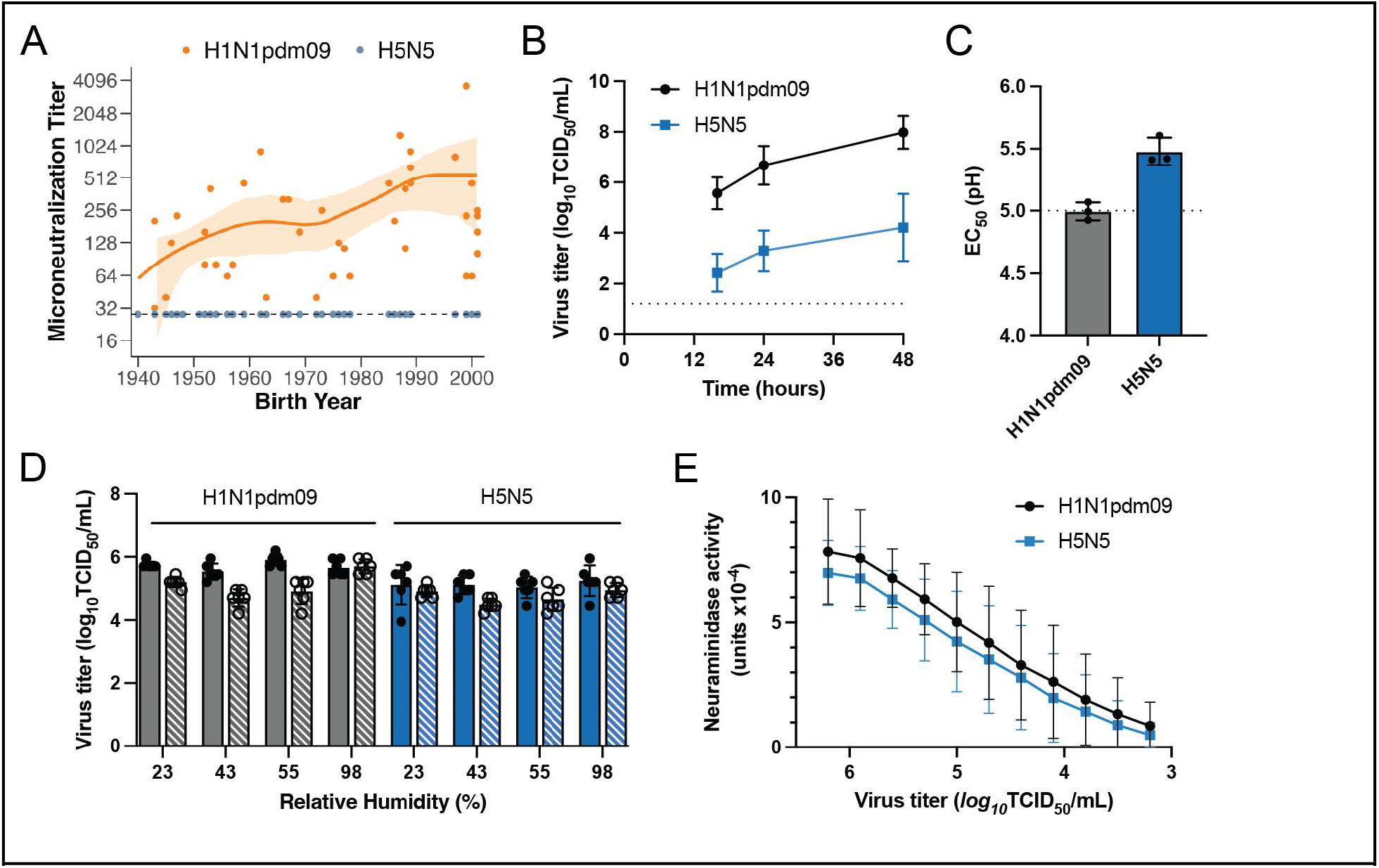
H5N5 displays in vitro characteristics that are a moderate risk. **A**. Sera from individuals collected in Fall 2020 from Pittsburgh, Pennsylvania were tested for neutralizing antibodies against A/California/07/2009 (H1N1pdm09, black circles) and A/Washington/2148/2025 (H5N5, blue circles) by microneutralization assay. The dashed line indicates the limit of detection. Trend lines represent generalized additive model (GAM) fits with 95% confidence intervals (shaded region). **B**. Transwells of HBE cell cultures grown at an air-liquid interface from three different patient donors were infected in triplicate with 10^3^ TCID_50_/mL of H1N1pdm09 or H5N5, and virus was collected at 16-, 24- and 48-hours post infection for titration on MDCK cells by TCID_50_ assay. Data are presented as the mean viral titer ± standard deviation. The dashed line indicates the limit of detection. All time points are significantly different p<0.0001. **C**. H1N1pdm09 and H5N5 were each incubated in a range of pH-adjusted PBS buffers for 1 hour before titering by TCID_50_ assay on MDCK cells. The data were fitted with an asymmetric sigmoidal curve and regression analysis of the dose-response curve was used to determine the EC_50_ which represents the pH of inactivation. The mean ± standard deviation corresponds to three independent biological replicates, each performed in triplicate. **D**. H1N1pdm09 (gray) and H5N5 (blue) virus stocks were diluted 1:10 in airway surface liquid (ASL) collected from three different HBE cell cultures from panel B. Ten 1uL droplets were spotted onto plastic plates and incubated for 2 hours at the indicated relative humidity in a controlled desiccator. Samples were collected immediately after plating (time 0, solid bars) and after 2 hours (striped bars) and titered on MDCK cells. Data represent the mean values ± standard deviation from three independent replicates each using ASL from a different HBE patient cell culture performed in duplicate. **E**. Neuraminidase activity for H1N1pdm09 and H5N5 was determined using an enzyme-linked lectin assay (ELLA) with fetuin as a substrate. The data are displayed as the mean ± standard deviation of four independent assays performed in duplicate.

### Molecular characterization of H5N5 demonstrates few pandemic risk features

Several molecular features are associated with the epidemiological success of IAV in humans including efficient replication in human respiratory cells, pH of inactivation similar ≤5.0, highly active neuraminidase (NA), and stable environmental persistence (9–14). Human bronchiole epithelial (HBE) cultures were used to assess viral replication, with H1N1pdm09 producing almost twice as much virus as H5N5 at all time points tested (Figure 1B), indicating that H5N5 is not well adapted to replicate in human airway epithelial cells. Viral entry necessitates that hemagglutinin (HA) undergoes a conformational change upon acidification in the endosome. The pH of inactivation for H5N5 was determined to be 5.5 (Figure 1C), which is higher than that of H1N1pdm09 with a pH of inactivation of 5.0. Prolonged IAV environmental persistence at mid-range humidity conditions is important for transmission (15), H5N5 was found to be as resistant as H1N1pdm09, to RH-mediated decay across all the conditions tested (Figure 1D), suggesting that H5N5 is stable in the environment. Finally, NA activity is essential for virion release from infected cells and higher NA activity has been linked with airborne transmission (12, 16). An enzyme-linked lectin assay indicated that H5N5 and H1N1pdm09 display similar NA activity across the entire dilution series (Figure 1E). These results indicate that H5N5 is resistant to environmental decay and has an active NA but is unable to replicate efficiently in HBE cells and has a high pH of inactivation. Thus, this lethal human H5N5 strain has only a few molecular characteristics thought to contribute to the epidemiological success of influenza viruses in humans, but these may be sufficient for transmission via close contact.

### No direct contact transmission in ferrets with no prior immunity

To address the potential risk of onward transmission of this H5N5 strain, H5N5 infected donor ferrets were cohoused with immunologically naïve recipients (Figure 2A). Only two of three donors shed virus in their nasal washes during the direct contact exposure period (Figure 2B). No infectious virus was detected in any recipient nasal wash, and none seroconverted to H5N5 at day 14 post exposure, demonstrating that this H5N5 strain does not transmit by direct contact under these study conditions.

**Figure 2.**
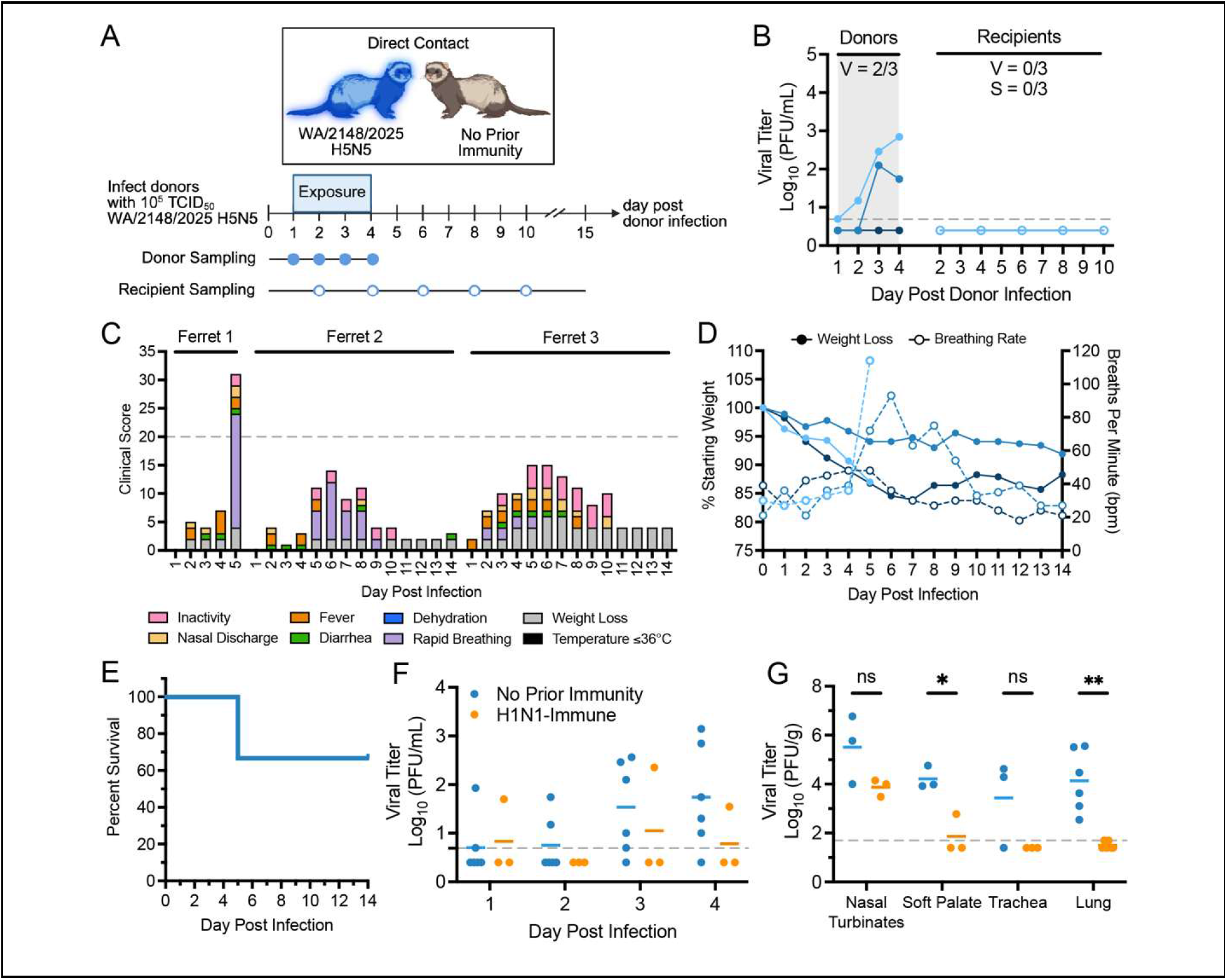
H5N5 does not transmit by direct contact but has moderate pathogenesis. Recipient ferrets with no prior immunity were cohoused with donor ferrets infected with 10^5^ TCID_50_ of H5N5 at one day post infection for the next 3 days. Nasal washes and oral swabs were collected from donors and recipients at indicated time points (circles). **B**. Viral shedding in nasal washes from donors and recipients was measured by TCID_50_ assay performed on MDCK cells. The number of ferrets that shed infectious virus (V) or seroconverted (S) as determined hemagglutination inhibition assay are indicated as fractions. The gray box indicates the exposure window. C. Survival was assessed in ferrets with no prior immunity infected with H5N5. Clinical signs were recorded. Humane endpoint was reached when clinical score was ≥20. **D**. Weight loss (left y-axis, closed circles) and breathing rate (right y-axis, open circles) were compared for survival study ferrets from panel C. Ferrets were euthanized at 22% weight loss or when breathing rate was >100 breaths per minute (bpm). **E**. Survival curve of ferrets with no prior immunity infected with H5N5 in panel C. **F**. Ferrets were infected with 10^5^ TCID_50_ of H1N1pdm09 three months prior to the H5N5 challenge. Ferrets with no prior immunity (N=6, blue) or with H1N1pdm09 prior immunity (N=3, orange) were intranasally inoculated with 10^5^ TCID_50_ of H5N5. Viral shedding in nasal washes from donors and recipients was measured by TCID_50_ assay on MDCK cells. The mean is indicated by the solid line. **G**. Groups of 3 H5N5 infected ferrets with no prior immunity (blue) or with H1N1pdm09 prior immunity (orange) were euthanized at 4 days post infection to measure viral load in respiratory tissues. Two lung lobes were analyzed separately. The mean is indicated by the solid line. Statistical analysis measured by unpaired t-test with Holm-Šídák correction. ^*^p<0.05; ^**^p<0.01.

To assess H5N5 pathogenicity, naïve ferrets were infected with H5N5 and were monitored for 14 days post infection. One ferret was euthanized at 5 days post infection, upon reaching humane endpoint, due to a very high breathing rate (>100 bpm vs baseline ≤39 bpm) (Figure 2C-D), resulting in a 33% (1/3) mortality rate (Figure 2E). While no neurological signs or temperature decline was observed, all ferrets displayed clinical signs of infection such as lethargy, weight loss, fevers and rapid breathing rates (Figure 2C).

### Prior immunity to H1N1pdm09 reduces H5N5 viral load and clinical disease

We have previously shown that ferrets with pre-existing H1N1pdm09 immunity are protected against severe disease induced by 2.3.4.4b H5N1 HPAI strains (17, 18). To determine whether pre-existing H1N1pdm09 immunity would protect against H5N5 pathogenesis, ferrets with no prior immunity or with pre-existing H1N1pdm09 immunity were intranasally infected with 10^5^ TCID_50_ of H5N5. Prior H1N1pdm09-immunity did not significantly impact viral shedding in nasal washes over the first four days of infection, although there was a trend toward lower viral load in the animals with prior immunity (Figure 2F). Importantly, on day 4 post infection, H1N1pdm09-immune ferrets had lower viral load in respiratory tissues compared to ferrets with no prior immunity, particularly in the soft palate and lungs (Figure 2G). H1N1pdm09-immune ferrets also displayed fewer clinical signs compared to ferrets with no prior immunity, consistent with reduced disease (data not shown). Overall, this data shows that pre-existing H1N1pdm09 immunity reduces H5N5 pathogenesis, indicating that H5N5 presents a low risk to a population with prior immunity.

## Discussion

The H5N5 (A/Washington/2148/2025) virus that caused a lethal human infection in 2025 was characterized for its pandemic potential with *in vitro* characterization of H5N5 revealing no neutralizing antibodies across a panel of human sera, indicating widespread susceptibility in the population. However, *in vivo* assessment indicated that prior H1N1pdm09 immunity reduced viral burden and disease signs during H5N5 infection, suggesting that there is some protection in the human population from prior IAV infection. This observation is of particular interest given evidence that cross-reactive NA immunity to N1 from human seasonal H1N1 strains was protective against H5N1 infections (19–21), suggesting that there is likely another mechanism of action since the NA proteins are distinct subtypes.

In contrast to our data, characterization of two closely related (>99% nucleotide identity) H5N5 strains by Erdelyan et al. demonstrated high mortality in ferrets after infection (6). These two representative strains were isolated from a bird (crow) and a mammal (raccoon) and displayed a preference for binding avian influenza virus receptor, 3’SLN-linked sialic acid (6). Sequence comparison of the crow and racoon strains as well as the lethal human H5N5 isolate characterized herein indicates that there are no differences at canonical mammalian adaptive sites (Supplemental Table 1). The raccoon strain had modest transmission to two of three cohoused recipient ferrets, with one displaying clinical disease and another one only seroconverting, whereas we did not observe transmission to our cohoused recipients (Figure 2B). Differences between the study outcomes could be due to the dose of the crow and racoon inoculums, which was 10x higher, differences in virus strains, and the duration of the cohoused exposure. A limitation of our study includes the assessment of only one H5N5 isolate, but there are currently no other human H5N5 isolate to compare to. Further characterization of circulating H5N5 strains is warranted to identify potential strains of higher pandemic potential.

H5N5 continues to circulate in wild bird populations in the US and Europe (22, 23) and thus will continue to evolve and adapt to new hosts. Ongoing surveillance and characterization studies are therefore warranted given its prevalence. Our *in vitro* studies provide guidance for targeted risk assessment studies in the future. We observed that H5N5 was stable in the environment at mid-range humidity levels and had NA activity similar to the H1N1pdm09 strain. However, H5N5 pH of inactivation was higher than 5.0, and this virus had poor replication capacity in HBE cultures. Rather than full characterizations of evolving H5N5 strains, future studies could focus on pH of inactivation and HBE growth. Emerging isolates with higher replication fitness in HBE cultures and pH of inactivation ≤5.0 will be of higher risk and could then be tested for ferret transmission.

## Acknowledgements

This project has been funded in part with Federal funds from the National Institute of Allergy and Infectious Diseases, National Institutes of Health, Department of Health and Human Services, under Contract No. 75N93021C00015; NIH award (UC7AI180311) from the National Institute of Allergy and Infectious Diseases (NIAID) supporting the Operations of The University of Pittsburgh Regional Biocontainment Laboratory (RBL) within the Center for Vaccine Research (CVR). We thank Dr. Rachel Duron for critical review and feedback.

## Author contributions

Conceptualization: V.L., S.S.L.

Formal analysis: M.N.V., G.E.Q., V.L., S.S.L. Funding acquisition: A.K.M., V.L., S.S.L.

Investigation: M.N.V., G.E.Q., A.E.S., K.B.-T., A.L.D., K.R.D., G.H., V.L.

Project administration: V.L., S.S.L. Resources: J.S., A.K.M.

Supervision: V.L., S.S.L. Visualization: V.L., M.N.V, G.E.Q.

Writing - original draft: M.N.V., G.E.Q., V.L., S.S.L.

Writing - review & editing: M.N.V., G.E.Q., A.K.M., V.L., S.S.L.

## Data availability

All raw data and sequence information uploaded to FigShare (10.6084/m9.figshare.32987585).

## Methods

### Human subjects research ethics statement

The University of Pittsburgh Institutional Review Board approved protocol STUDY20030228 for collection of human serum samples from healthy adult donors who provided written informed consent for their samples to be used in infectious disease research. All participants self-reported their age, sex, race, ethnicity, residential zip code, history of travel and immunization. Human bronchiole epithelial cell cultures were obtained from deidentified patients under the University of Pittsburgh Institutional Review Board approved protocol STUDY19100326.

### Animal research ethics statement

All ferret experiments were conducted in Emory University’s BSL3 facility in compliance with the guidelines of the Institutional Animal Care and Use Committee (Emory University’s approved protocol 2024-00000112). Isoflurane was used to sedate animals for all nasal washes and oral swabs, as directed by approved methods. For terminal procedures, animals received ketamine and xylazine for sedation, followed by euthanasia solution administered via cardiac injection.

### Cells

Madin-Darby canine kidney (MDCK) cells and 293T cells were obtained from American Type Culture Collection (ATCC) and maintained in Minimum Essential medium. Medium was supplemented with 10% fetal bovine serum, 2 mM L-glutamine and 2 mM penicillin/streptomycin. Primary human bronchiole epithelial (HBE) cell cultures were differentiated from human lung tissue and cultured at an air-liquid interface using a protocol approved by the institutional review board at the University of Pittsburgh. All cells were incubated at 37^°^C with 5% CO_2_.

### Virus Rescue

The reverse genetic plasmids of A/California/07/2009 (H1N1pdm09) were a generous gift from Dr. Jesse Bloom (Fred Hutch Cancer Research Center, Seattle). Reverse genetics plasmids expressing A/Washington/2148/2025 were synthesized based on sequence deposited in GISAID (Accession number EPI_ISL_20252012). Noncoding regions (NCRs) were based on a set of pan-avian H5Nx assembled trees to identify the most conserved sequences across H5Nx strains. Sequences synthesized into reverse genetics plasmids are available in the data repository listed in Data Availability section. The eight reverse genetics plasmids were transfected into 293T cells using Lipofectamine 2000 in Opti-MEM complete media. Twenty-four hours later, supernatant from transfected 293T cells were overlayed on top of MDCK cells with MEM supplemented with 4 mM L-glutamine, 1x antibiotic-antimycotic and 1 μg/mL of TPCK-treated trypsin. The 293T supernatant overlay on top of MDCK cells was repeated at 48 hours post transfection. MDCK cells were monitored for cytopathic effect (CPE) for 2 days post final supernatant overlay, and supernatant was collected (cell passage 1, cP1). A second cell passage (cP2) stock was generated on MDCK cells and used for all subsequent experiments. Plasmids were sequence verified prior to rescue.

### Virus titration

For TCID_50_ virus titration, ten-fold serial dilutions were made with 20 μL of sample being diluted into 180 μL of Minimum Essential medium (supplemented with 2 mM L-glutamine and 1X Anti-Anti) in 96-well plates of confluent MDCK cells and performed in quadruplicate. Cytopathic effect (CPE) was determined at 4 dpi. Virus titers were calculated using Reed and Muench method (24) and expressed as log_10_ TCID_50_/mL. The limit of detection for this assay is 10^1.2^ TCID_50_/mL.

### Microneutralization assay

For this assay, human sera was treated with receptor destroying enzyme (RDE) overnight at 37^°^C (3 volumes of 2X RDE to 1 volume of sera). The following day, the sera was heat inactivated at 56^°^C for 30 minutes and 6 volumes of saline were added for a final dilution of 1:10. Two-fold serial dilutions of RDE-treated human serum in a volume of 125 mL were incubated with 125 uL of 10^3.3^ TCID_50_ of A/California/07/2009 or A/Washington/2148/2025 H5N5 for 1 hour at room temperature with continuous rocking. 150 uL of Minimum Essential medium supplemented with 2 mM L-glutamine and 1X Anti-Anti with TPCK was added to 96-well plates with confluent MDCKs before 50 uL of virus:serum mixture was added in quadruplicate. After 4 days, CPE was determined and the neutralizing antibody titer was expressed as the reciprocal of the highest dilution of serum required to completely neutralize the infectivity of each virus on MDCK cells. The concentration of antibody required to neutralize 100 TCID_50_ of virus was calculated based on the neutralizing titer dilution divided by the initial dilution factor, multiplied by the antibody concentration. The limit of detection for this assay is 28.

### Replication kinetics in human bronchiole epithelial cell cultures

Three different HBE patient cell cultures were used (HBE0527, HBE0530, HBE0531) with triplicate transwells for each time point. The apical surface of the uninfected HBE transwells was washed in 150 uL phosphate-buffered saline (PBS) to remove any airway surface liquid for 10 minutes and collected for use in stability experiments. An inoculum of 10^3^ TCID_50_ of A/California/07/2009 or A/Washington/2148/2025 H5N5 was added per 100 uL of HBE growth medium. The titer of each inoculum was determined to ensure that both strains were within 10^0.5^ TCID_50_/mL of each other. After 1 hour incubation, the inoculum was removed and the apical surface was washed three times with 150 uL of PBS. At the indicated time points, 150 uL of HBE medium was added to the apical surface for 10 minutes to capture released virus particles. Infectious virus was quantified by TCID_50_ assay using the endpoint method. The limit of detection for this assay is 10^1.2^ TCID_50_/mL.

### Stability of stationary droplets

Airway surface liquid (ASL) was collected from uninfected HBE patient lines HBE0527, HBE0530, HBE0531 using 150 uL of PBS per transwell for 10 minutes at 37^°^C. Virus stock was mixed with ASL at a ratio of 1:10 and used to generate ten 1 uL droplets in triplicate on a 6-well plate with tissue culture-treated plastic (ThermoFisher, Waltham, MA). Virus:ASL droplets were incubated in a sealed desiccator chamber containing saturated salt solutions of potassium acetate, potassium carbonate, magnesium nitrate and potassium sulfate to maintain desired 23%, 43%, 55% and 98% relative humidity (RH), respectively. Chambers were maintained in a biosafety cabinet, and a HOBO UX100011 data logger (Onset, Cape Cod, MA) was placed in the desiccator to collect RH and temperature data. After 2 hours, the droplets were collected in 500 uL of L-15 Leibovitz’s medium (Gibco, Grand Island, NY), which was titered on MDCK cells using the TCID_50_endpoint method (24).

### Ferret screening

Prior to purchase from Triple F Farms (Sayre, PA), sera from six-to eight-month-old male ferrets were screened for antibodies against influenza A and B viruses using hemagglutination inhibition (HAI). Antigens used for screening were obtained through the International Reagent Resource, Influenza Division, WHO Collaborating Center for Surveillance, Epidemiology and Control of Influenza, Centers for Disease Control and Prevention, Atlanta, GA, USA: CDC Antigen, Influenza A(H1N1)pdm09 Control Antigen (A/Victoria/2570/2019), BPL-Inactivated, FR-1820; CDC Antigen, Influenza A(H3) Control Antigen (A/Delaware/01/2021), BPL-Inactivated, FR-1821; CDC Antigen, Influenza B Control Antigen, Victoria Lineage (B/Michigan/01/2021), BPL-Inactivated, FR-1822.

### Hemagglutinin inhibition (HAI) assay

RDE-treated sera were serially diluted two-fold and incubated with eight hemagglutinating units of HA antigen or virus. After 15 minutes of incubation, an equal volume of 0.75% turkey red blood cells (Lampire Biological Laboratories) was added and incubated for 30 minutes. The reciprocal of the highest dilution of serum that inhibited hemagglutination was determined to be the HAI titer. The limit of detection for this assay is 1:10.

### Transmission study

Donor ferrets were intranasally inoculated with 10^5^ TCID_50_ of A/Washington/2148/2025 in 500 μl. At day 1 post donor inoculation, donors and recipients with no prior immunity were paired and cohoused for 3 days of direct-contact exposure. Pairs were separated and single-housed after exposure. Nasal washes and oral swabs were collected from donors for 4 days post inoculation before euthanasia at predetermined time points. Nasal washes and oral swabs were collected from recipients for 10 days post exposure or until reaching humane endpoint criteria. Sera was collected from recipients at 14 days post exposure or upon reaching humane endpoint and used in HAI assays to determine seroconversion. Recipients were euthanized at 14 days post exposure if not euthanized early due to reaching humane endpoint criteria.

### Pathogenesis and survival in ferrets

To generate ferrets with prior H1N1pdm09-immunity, ferrets were intranasally inoculated with 10^6^ TCID_50_ in 500 uL of A/California/07/2009 H1N1pdm09 and their immunity was allowed to wane for 3 months prior to infection with H5N5. Ferrets with no prior immunity or with prior H1N1pdm09-immunity were then intranasally inoculated with 10^5^ TCID_50_ of A/Washington/2148/2025 in 500 μl. Nasal washes and oral swabs were collected from ferrets for 4 days post inoculation. Ferrets were euthanized at day 4 post inoculation to measure viral load in respiratory tissues. A subset of inoculated ferrets with no prior immunity were monitored past day 4 post inoculation to assess survival and were euthanized upon reaching humane endpoint.

### Ferret clinical signs

Ferrets were monitored for signs of clinical disease throughout the study to assess disease severity with scoring expanded upon what was previously described (25). Humane endpoint is met at a cumulative clinical score ≥20. Clinical signs were assigned point values as follows: Activity (0 = alert and playful; 2 = alert but playful only when stimulated; 4 = alert but not playful when stimulated; 10 = neither alert nor playful when stimulated; 20 = moribund), Temperature (0 = normal 36-40°C; 2 = fever ≥40°C; 4 = severe fever ≥40°C + 2°C more than baseline; 20 = hypothermia <36°C), Weight Loss (0 = 0-5%; 2 = 5-10%; 4 = 10-15%; 6 = 15-20%;10 = 20-22%; 20 = ≥22%), Nasal Discharge (0 = none; 1 = mild; 2 = moderate; 3 = severe),Respiratory Rate (≤39 breaths per minute [bpm] = 0; 42-57 bpm = 2; 60-87 bpm = 5; 90-99 bpm = 10; >100 bpm = 20), Dehydration (0 = none; 3 = skin tent ≥1 seconds), and Diarrhea (0 = none; 1 = present).

